# Mosquito defensins enhance Japanese encephalitis virus infection by facilitating virus adsorption and entry within mosquito

**DOI:** 10.1101/2020.04.28.065904

**Authors:** Ke Liu, Changguang Xiao, Shumin Xi, Muddassar hameed, Abdul Wahaab, Donghua Shao, Zongjie Li, Beibei Li, Jianchao Wei, Yafeng Qiu, Denian Miao, Huaimin Zhu, Zhiyong Ma

## Abstract

Japanese encephalitis virus (JEV) is a viral zoonosis which can cause viral encephalitis, death and disability. *Culex* is the main vector of JEV, but little is known about JEV transmission by this kind of mosquito. Here, we found that mosquito defensin facilitated the adsorption of JEV on target cells via both direct and indirect pathways. Mosquito defensin bound the ED III domain of viral E protein and directly mediated efficient virus adsorption on the target cell surface, Lipoprotein receptor-related protein 2 expressed on the cell surface is the receptor affecting defensin dependent adsorption. Mosquito defensin also indirectly down-regulated the expression of an antiviral protein, HSC70B. As a result, mosquitos defensin enhances JEV infection in salivary gland while increasing the possibility of viral transmission by mosquito. These findings demonstrate that the novel effects of mosquito defensin in JEV infection and the mechanisms through which the virus exploits mosquito defensin for infection and transmission.

## Introduction

Japanese encephalitis virus (JEV), a member of *Flaviviridae flavivirus*, is prevalent in Asia-Pacific tropical and subtropical regions [1–3]. JEV is mainly transmitted through mosquito bites [2, 4]. Pigs are reservoir hosts for JEV, and humans, horses and other animals are dead-end hosts [2, 5]. Because the prevention and control of JEV rely on vaccines with a limited window of protection [6–8], JEV can easily cause death or permanent disability. More than 100,000 people are at risk of JEV infection, and immunocompromised children and older individuals are at particular risk [9, 10]. The World Health Organization has reported that more than 67,900 cases of JEV infection globally each year, more than 10,000 of which are fatal. As global temperature increases, the clinical incidence of Japanese encephalitis is increasing as well, owing to an increase in the habitat range and activity of mosquitoes carrying JEV as the climate warms [2, 4, 9]. Few studies have addressed the transmission mechanism of JEV by mosquito vectors [4]. Thus, a detailed understanding of the interaction between JEV and mosquito vectors will be essential to improve control of JEV transmission.

*Culex* is the principal vector of JEV [11, 12]. The virus can spread throughout the mosquito body, including salivary glands [13]. When an infected mosquito bites humans or animals, the virus is transmitted to the skin through the saliva. The mosquito vector also induces an immune response to JEV [14–16]. For example, C-type lectin and a series of proteins increase rapidly after infection [17, 18]. C-type lectin plays an important role in infection by JEV and other *Flaviviridae* viruses in mosquitoes, but the role of defensin has not yet been clearly characterized.

Defensins are antimicrobial peptides consisting of 25–60 amino acids that are produced by innate immune system [15, 19]. Defensin is one of the crucial immune effectors in insects [20]. The antiviral effects of defensins have been well described in mammalian cells. Human defensins have been reported to inhibit herpes simplex virus type 2 (retrocyclin-1, retrocyclin-2) [21], human immunodeficiency virus (human beta defensin-1, human beta defensin-2, Human beta defensin-3) [22, 23] and other viruses. However, human beta defensin-6, expressed by adenovirus vectors, enhances parainfluenza virus type 3 replication [24]. Normally, mammalian defensins can directly destroy the virus particles by binding to the surface of envelope protein. They can also interact to the cell surface receptor and influence cell signal transduction [19, 25]. Although there are many differences between the mammalian and mosquito immune systems, defensins are considered important effectors in the mosquito immune response. Therefore, the role of mosquito defensins during the process of JEV infection requires further study.

In this study, we observed complex roles of mosquito defensin in JEV infection: a weak antiviral effect and a strong effect enhancing binding. In the latter, defensin directly binds the ED III domain of the viral E protein and promotes the adsorption of JEV to target cells by interacting with lipoprotein receptor-related protein 2 (LRP2), thus accelerating virus entry. Mosquito defensin also down-regulates the expression of the antiviral protein HSC70B on the cell surface, thus facilitating virus adsorption. Together, our results indicate that the facilitation effect of mosquito defensin plays an important role in JEV infection and potential transmission.

## Results

### JEV infection up-regulates defensin expression *in vivo* and *in vitro*

Defensin is one of the major innate immunity effectors in mosquitoes. To study the role of defensin in JEV infection, we first assessed the expression of defensin in *Culex pipiens pallens* (*Cpp*) which is the natural vector of JEV after infection. Five-day-old female mosquitoes after emergence were infected by a microinjector with a dose of 1000 MID_50_ [18]. The mosquitoes were collected 4, 7 and 10 days after injection, and the JEV E mRNA levels in the whole body, salivary gland and midgut were determined. JEV E mRNA showed higher levels in the salivary gland than in the whole body and midgut (Fig. 1A). At 10 days, the JEV E mRNA level increased dramatically, thus indicating that the virus reproduced rapidly during this period. For instance, JEV E mRNA levels increased by 9.7- and 11.7-fold at 10 days compared with 7 days in the whole body and midgut, respectively. A greater increase was observed in the salivary gland, reaching 14.9 fold at 10 days compared with 7 days. Because high virus levels in mosquitoes were observed 7 and 10 days post JEV infection, we then determined the defensin A and total defensins mRNA level in the whole body on 7 and 10 days. *Cpp* defensin A mRNA levels on both days were significantly higher in the JEV infection group than the control group, although the level decreased slightly at 10 days (Fig. 1B). Change levels of total defensins showed the similar trend (Fig. 1B). Furthermore, we compared *cpp* defensin A and total defensins mRNA levels in the whole body, salivary gland and midgut. Defensin A and total defensins mRNA showed similar levels of up-regulation in the salivary gland and whole body, which were higher than those in the midgut (Fig. 1C). Significantly higher mRNA levels (*p*<0.005) were observed in the JEV infection group than the control group for whole body, salivary and midgut. This suggested that *Cpp* defensin A expression was positive correlated with JEV infection in mosquitoes.

**Fig. 1.**
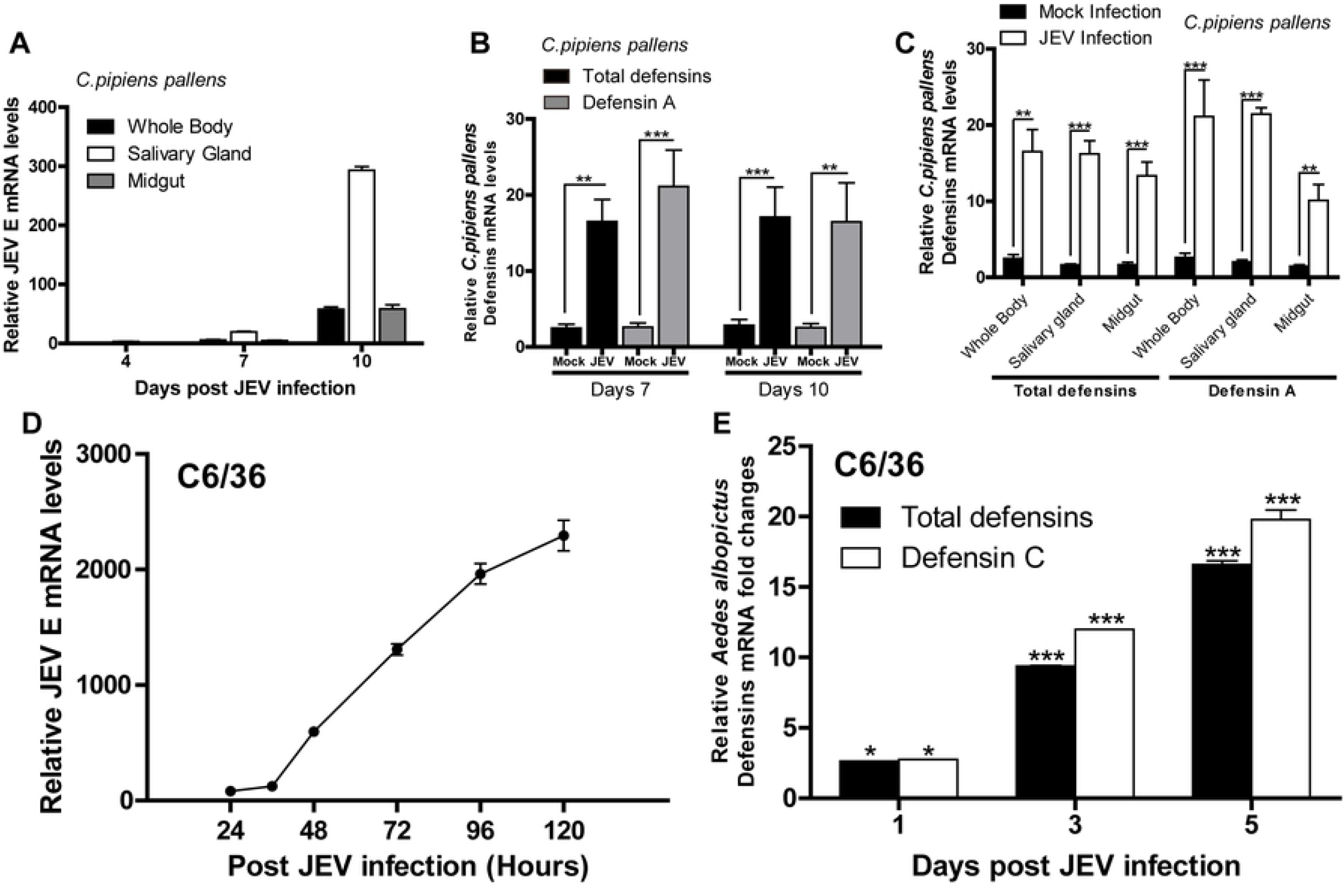
JEV infection induced defensin up-regulation in mosquito vectors. (A) JEV infection curve in mosquitoes. JEV (10^3^ MID_50_) or PBS was inoculated into female mosquitoes by throat injection. Whole body, salivary gland and midgut samples were collected at 4, 7 and 10 days post JEV infection. JEV E expression was quantified by real-time PCR. (B) Expression levels of *cpp* defensins in the whole body. JEV (10^3^ MID_50_) or PBS was inoculated into female mosquitoes by throat injection. *Cpp* defensin A mRNA levels in the whole body at 7 and 10 days post JEV infection were quantified by real-time PCR. (C) Expression levels of *cpp* defensins in the midgut and salivary gland. JEV (10^3^ MID_50_) or PBS was inoculated into female mosquitoes by throat injection. Midgut and salivary gland were separated at 7 days post infection. *Cpp* defensin A mRNA levels were quantified by real-time PCR. (D) One step growth curve of JEV virus in C6/36. C6/36 cells were infected with 5 MOI and collected at different time points, as shown in Fig. 1D. JEV E mRNA levels were quantified by real-time PCR. (E) Expression levels of *aa* defensins. C6/36 cells were infected with 5 MOI and collected at 1, 3 and 5 days post JEV infection. *Aa* defensin C and total defensins mRNA levels were quantified by real-time PCR. All experiments were done in triplicate and were performed at least three times. Data are shown as Mean values ± standard deviations.

The gene encoding *cpp* defensin was not found in the NCBI database. According to PCR amplification (Fig. S1A), sequencing and BLASTn (https://blast.ncbi.nlm.nih.gov) results, we found two gene types of defensin: defensin A (submitted to NCBI with accession number MH756645) and an incomplete information defensin. The mature protein regions of these two defensins shared 99.5% sequence similarity (Fig. S1B). We designed specific primers (Table. S1, Fig. S2) for real-time PCR detection according to the *Cpp* defensin A sequence. However, no specific primers were available for the unnamed defensin, because scarce specific sequence was obtained. Therefore, we quantified the mRNA copy number of total and type A defensins to determine which subtype is the primary defensin in *cpp* (Fig. S1C). The fold change of mRNA levels of defensin A were significantly higher (7 days, *p*<0.05) than or similar (10 days) to the total defensin levels (Fig. 1B), thus it implied that defensin A accounts for the majority of total defensins. To analyze whether defensin functioned universally among organisms, we aligned the defensin protein sequences of mosquito vectors of flavivirus. We also quantified the mRNA copy number of total and type C defensins to determine which subtype is the primary defensin in *Aedes albopictus* (*aa*) (Fig. S1C). *Aedes* defensin C is the major type of defensin in C6/36 cell, a cell line from *Aedes albopictus*. The sequence similarities were all above 97.6% between mosquito vectors (Fig. S1D), suggesting that mosquito defensins serve similar functions. In contrast, the sequence similarities were significantly low between mosquitoes and human (Fig. S1E). We then used *cpp* defensin A (accession number MH756645) and *aa* defensin C (accession number XP_019527114.1) to study the functions of defensin in JEV infection within mosquito vectors.

To confirm the up-regulation of defensin in different mosquito vectors caused by JEV infection, we infected C6/36 cells with JEV *in vitro*. JEV E mRNA levels increased from 24 h to 120 h post JEV infection (Fig. 1D). We further analyzed the change of total defensins and primary defensin (*aa* defensin C, Fig. S1C) after JEV infection. *aa* defensin C mRNA levels were up-regulated to 2.75-, 11.9-, 19.7-fold in JEV infection compared with mock infection at 1, 3 and 5 days, respectively (Fig. 1E). Also total defensins were up-regulated after JEV infection (Fig. 1E). Together, our results indicated that defensin levels were up-regulated after both *in vivo* and *in vitro* infections.

### Mosquito defensin shows species specificity in facilitating JEV infection

Mature defensin is an extracellular protein which length is less than 60 amino acids. To confirm the function of defensin on JEV infection, we synthesized mature defensin peptides with high purity (⩾99%) to perform further analysis. Scrambled defensin peptides were used as controls. *Cpp* defensin A and *aa* defensin C peptides were used in both *in vivo* and *in vitro* experiment. Defensins and JEV were pre-mixed before injection into mosquitoes. Unexpectedly, in *in vivo* experiment, JEV E mRNA levels increased by 2.95- and 6.13-fold in the *cpp* defensin A treated groups compared with the control groups at 7 and 10 days post infection, respectively (Fig. 2A). And the same changing trend of JEV level was observed in *aa* defensin C treated group (Fig. 2A). We also confirmed this enhancement of defensins on JEV infection by RNA interference. siRNA sequences target *Cpp* defensins or *Cpp* defensins A were designed and used in *in vivo* RNA interference. JEV E mRNA levels were decreased by more than 5 fold in *Cpp* defensin siRNA groups and more than 3 fold in *Cpp* defensin A siRNA groups compared to scramble group (Fig. 2B, Fig. S3A). Indirect immunofluorescence assay (IFA) analysis also showed higher JEV E levels in the mosquito defensin treated cells than the control (Fig. 2C), and lower JEV E levels in the mosquito defensin knockdowned cells than the control (Fig. 2C).

**Fig. 2.**
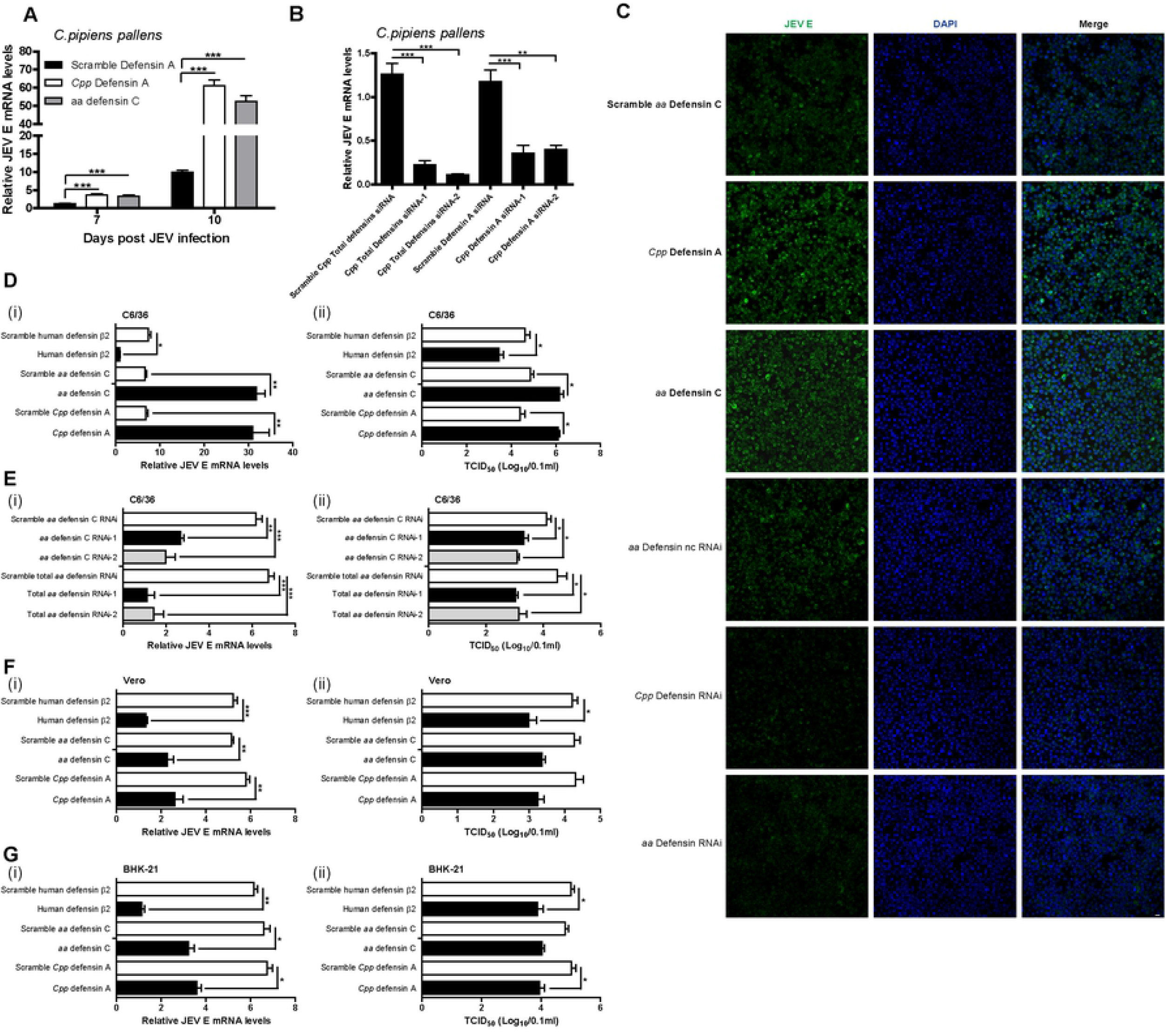
Mosquito defensin facilitated JEV infection in mosquito vectors. (A) Mosquito defensin facilitated JEV infection in *culex* mosquitoes. Mosquito defensins (100 μM) and JEV (10 MID_50_) were pre-mixed at 4°C for 2 h and inoculated into female mosquitoes. JEV E mRNA levels in the whole body at 7 and 10 days post JEV infection were quantified by real-time PCR. (B) Mosquito defensins interference harmed JEV infection in *culex* mosquitoes. Female mosquitoes were injected with siRNAs for 3 days, then JEV was injected in dose of 10 MID_50_. JEV E mRNA levels in the whole body at 6 days post JEV infection were quantified by real-time PCR. (C) Mosquito defensins facilitated JEV infection on C6/36 in IFA detection. Mosquito defensins (50 μM) and JEV (0.5 MOI) were pre-mixed at 4°C and inoculated into C6/36 cells for 2 h (upper three panels). siRNAs target defensin were transfected into C6/36 cell for 24 h, then JEV was inoculated to C6/36 cell for 2 h (lower three panels). IFA was performed on cells at 3 days post infection. Bar, 10 μm. (D) Mosquito Defensins facilitated JEV infection in C6/36 cells. Mosquito defensins (50 μM) and JEV (0.5 MOI) were pre-mixed at 4°C and inoculated into C6/36 cells for 2 h. The cells and supernatant were collected at 3 days post infection to quantify JEV E mRNA levels (i) and TCID_50_ (ii). (E) Defensin knockdown harmed JEV infection in C6/36 cell. siRNAs target defensin were transfected into C6/36 cell for 24 h, then equal JEV was added to C6/36 cell without changing media. The cells and supernatant were collected at 2 days post infection to quantify JEV E mRNA levels (i) and TCID_50_ (ii). (F - G) Mosquito defensins inhibited JEV infection in mammalian cells. Mosquito defensins (50 μM) and JEV (0.5 MOI) were pre-mixed at 4°C and were inoculated into Vero (F) or BHK (G) for 1.5 h at 37°C. The cells and supernatant were collected at 48 h post infection to quantify JEV E mRNA levels (i) and TCID_50_ (ii). All experiments were done in triplicate and were performed at least three times. Data are shown as Mean values ± standard deviations.

To compare the role of defensin from different species, human defensin β2 showed high antiviral activity was synthesized [25, 26]. Firstly, we compared the effects of *aa* defensin C, *Cpp* defensin A and human defensin β2 on C6/36 cells. *aa* defensin C enhanced JEV infection on C6/36, as indicated by both JEV E mRNA (4.88 fold) and TCID_50_ (1.3 titer) levels (Fig. 2D i and ii). Treating with *Cpp* defensing A also resulted in the enhancement of JEV infection. In contrast, human defensin β2 inhibited JEV replication on C6/36 cell, thus demonstrating that defensins from different species have diverse functions in JEV infection (Fig. 2D i and ii). To confirm this effect of mosquito defensin, we used siRNAs target defensin of C6/36. JEV was inoculated and detected after siRNAs transfection. JEV E mRNA levels decreased by 4.7 to 6 folds in *aa* defensin interference groups, and decreased by 2.3 to 3.1 folds in *aa* defensin C interference groups respectively (Fig. 2E i and ii, Fig. S3B). These results were consistent with the *in vivo* data.

To obtain detailed insight into the function of mosquito defensin, we analyzed the effects of mosquito defensin on mammalian cells contain Vero and BHK-21. *aa* defensin C reduced the JEV replication by 2.2 to 2.7 folds (Fig. 2F i and Fig. 2G i) and decreased JEV TCID_50_ levels by 0.6 to 0.8 titers (Fig. 2F ii and Fig. 2G ii), thus indicating that it inhibits JEV infection in mammalian cells as human defensin dose. Although the inhibition ability of *aa* defensin C was lower than that of human defensin β2, it still inhibited JEV replication. Therefore, the facilitation effects of mosquito defensin on JEV were valid only on mosquitoes and mosquito cells. To confirm that the effect of defensins was not due to cytotoxicity, we measured the IC_50_ of each defensin through MTT assays. The results showed that defensins had no significant cytotoxicity on cells (Table. S2).

### Mosquito defensin enhances JEV adsorption to target cells

To study the exact mechanisms of mosquito defensin facilitates JEV infection, we analyzed the influence of *aa* defensin C on different infection steps on C6/36 cell. As infection steps can be measured by temperature and time shift, we detected adsorption, uncoating and replication of JEV [27]. Adsorption was determined to be a key step in the facilitation effect (Fig 3A). Next, we detected JEV adsorption at different time points. JEV mixed with defensin or scrambled peptides was inoculated to C6/36 cells for different time points at 0°C. After being washed with PBS for three times, cells with absorbed JEV were collected. JEV E mRNA levels were determined by real-time PCR. C6/36 cells treated with *aa* defensin C showed significantly higher JEV E mRNA levels at 4 and 6 h post adsorption, proof that defensin enhanced JEV adsorption to C6/36 (Fig. 3B). IFA showed that JEV adsorption greatly increased in a time course of *aa* defensin C treatment (Fig. 3C and Fig. 3D). Both nuclear staining (DAPI) and membrane staining (Did) of C6/36 cell were conducted in IFA absorption analysis. There was stronger JEV adsorption in the *aa* defensin C groups than the control groups at each time points in both DAPI staining (Fig. 3C) and Did staining (Fig. 3D) cells. To study how mosquito defensin facilitated JEV adsorption, FITC labeled *aa* defensin C was used. Defensin-FITC and JEV were mixed before incubation at 0°C. After incubation, unabsorbed defensin and JEV were washed by PBS for five times. The cells were collected at the indicated time points to observe the co-localization of defensin and JEV. Strong co-localization on the cell surface was observed between *aa* defensin C and JEV (Fig. 3E) and increased over time. Thus, the facilitation effect of mosquito defensin on JEV was attributed to the binding between them. Additionally, JEV mixed with *cpp* defensin A showed high adsorption capacity in salivary glands (Fig. 3F). Take together, our results indicated that mosquito defensin is able to bind JEV and facilitate virus adsorption.

**Fig. 3.**
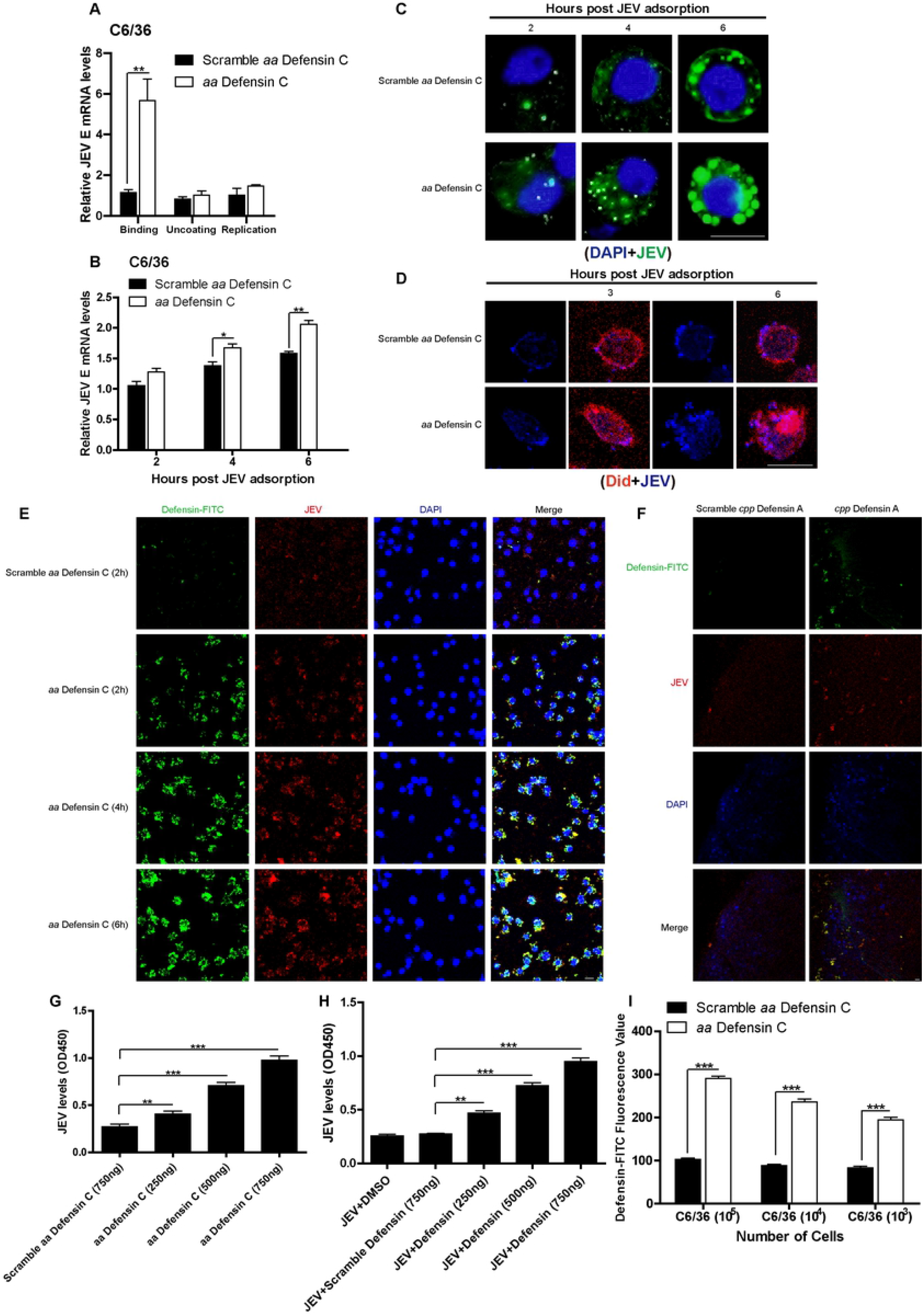
Mosquito Defensin facilitated JEV adsorption to mosquito cells. (A) Step scan of JEV infection on C6/36 cell. Steps of JEV infection on C6/36 cell were analyzed by different treatments. For binding, virus (0.5 MOI) and defensin (50 μM) were mixed and inoculated to C6/36 cell on ice for 4 h, washed with PBS for five times, cultured for 48 h with new media. For uncoating, virus (0.5 MOI) was inoculated to C6/36 cell on ice for 4 h, washed with PBS for five times. Fresh media with defensin (50 μM) was added to cell for 6 h. After incubation, cell was washed and cultured for another 42 h with new media. For replication, virus (0.5 MOI) was inoculated to C6/36 cell on ice for 4 h, washed with PBS for five times. New media without defensin was added to cell for 48 h, and defensin (50 μM) was added into media at 6 h post culture. The cells were collected to quantify JEV E mRNA levels by real-time PCR. (B) *Aa* defensin C facilitated JEV adsorption to C6/36 in a time-dependent manner. *Aa* defensin C (50 μM) and JEV (0.5 MOI) were pre-mixed at 4°C and inoculated into C6/36 cells on ice for 4 h. Unabsorbed JEV was removed by washing with PBS three times. The cells were collected to quantify JEV E mRNA levels by real-time PCR. (C and D) IFA assay of JEV adsorption to C6/36 cells. *Aa* defensin C-FITC (50 μM) and JEV (1 MOI) were pre-mixed at 4°C and inoculated into C6/36 cells on ice. Unabsorbed JEV and defensin were removed by washing with PBS three times. The cells were strained with antibody and DAPI (C) or Did (D). (E) Co-localization of defensin and JEV on the cell surface. *Aa* defensin C-FITC (50 μM) and JEV (1 MOI) were pre-mixed at 4°C and inoculated into C6/36 cells on ice for 2 h, 4 h and 6 h. Unabsorbed JEV and defensin were removed by washing with PBS three times. The cells were treated to observe JEV E (red fluorescence), defensin-FITC (green fluorescence) and nuclei (blue fluorescence). Bar, 10 μm. (F) Co-localization of *Cpp* defensin A-FITC and JEV on the salivary gland. The salivary glands from uninfected female mosquitoes were freshly isolated. Pre-mixed *cpp* defensin A-FITC (50 μM) and JEV (1 MOI) were added to salivary glands and incubated at room temperature for 1 h. JEV E was labeled with monoclonal antibody (red fluorescence). Defensin-FITC was detected by green fluorescence, and nuclei were stained with DAPI (blue fluorescence). Bar, 20 μm. (G) JEV bind to defensin. The plate was coated with *aa* defensin C, incubated with JEV and assessed with anti-JEV monoclonal antibody. (H) Defensin and JEV mixture binds to C6/36 cells. The plate after polylysine treatment was coated with C6/36, pre-mixed defensin and JEV were added, and detection was performed with anti-JEV antibody. (I) Defensin binds to C6/36 directly. The polylysine treated plate was coated with C6/36, defensin-FITC were added, and fluorescence value was detected. All experiments were done in triplicate and were performed at least three times. Data are shown as Mean values ± standard deviations.

The interaction between defensin and JEV was also confirmed by ELISA. The plate wells were coated with *aa* defensin C, incubated with JEV and next incubated with anti-JEV antibody. As expected, JEV bound defensin efficiently. Even with a 250 ng defensin coating treatment, the JEV level was significantly higher than that in the control group (Fig. 3G). To determine the adsorption capacity of the JEV-defensin complex to C6/36 cells, we coated plate wells with fresh C6/36 cells after polylysine treatment, added pre-mixed defensin and JEV, and detected JEV with anti-JEV monoclonal antibody. In accordance with the results of qPCR and IFA, the interaction of defensin with JEV significantly enhanced JEV adsorption to C6/36 cells (Fig. 3H).

Based on the previous results, we deduced that mosquito defensin can efficiently bind to cell surface. The interaction of mosquito defensin with the cell surface was assessed through ELISA. Defensin directly interacted with C6/36, and a higher FITC value than that of the control was observed (Fig. 3I). This finding implied that the facilitation effect of defensin on JEV was caused by increasing the affinity of JEV on the cell surface.

### Defensin directly binds the JEV ED III domain

Defensins can bind to viral envelope protein [25]. To precisely understand the interaction mechanisms of JEV enhancement mediated by defensin, we expressed the three structural proteins (C, prM and E) of JEV and the exposure area (ED III) of E protein through an S2 insect protein expression system [28, 29], and further purified these proteins via 6×His agarose. To analyze the interaction between viral proteins and defensins, two ELISA methods were used. The plate wells were coated with defensin, incubated with purified proteins and detected by corresponding antibodies. Scrambled defensin was used as control. Absorbance results showed high affinity between *aa* defensin C and E protein or ED III protein, which were ~0.95 and ~1.17, respectively (Fig. 4A). Consistent results were observed in defensin-FITC testing. The plate wells were coated with purified viral proteins and then incubated with *aa* defensin C-FITC or scrambled defensin-FITC. E and ED III proteins showed higher fluorescence values of 332 and 369, respectively (Fig. 4B). Both tests suggested that the ED III domain of E protein is the key region involved in *aa* defensin C and JEV binding. Subsequently, purified E and ED III proteins were mixed with *aa* defensin C and used to inoculate C6/36 cells at 0°C for 4 h. Unabsorbed defensin and JEV were removed by washing with PBS after incubation. The effect of *aa* defensin C on facilitating E and ED III adsorption was observed by fluorescence microscope (Fig. 4C and Fig. 4D). E proteins and ED III proteins bound more efficiently to the C6/36 cell with *aa* defensin. Additionally, *aa* defensin C showed co-localization with E or ED III protein on the C6/36 (Fig. 4C, merge panel). The same results were observed in membrane stained C6/36 cells. E proteins and ED III proteins bound more efficiently to the C6/36 cell surface with *aa* defensin, and *aa* defensin C also showed co-localization with E or ED III protein on the C6/36 surface (Fig. 4D). Thus indicating that the ED III domain of the JEV E protein responsible for binding with *aa* defensin C.

**Fig. 4.**
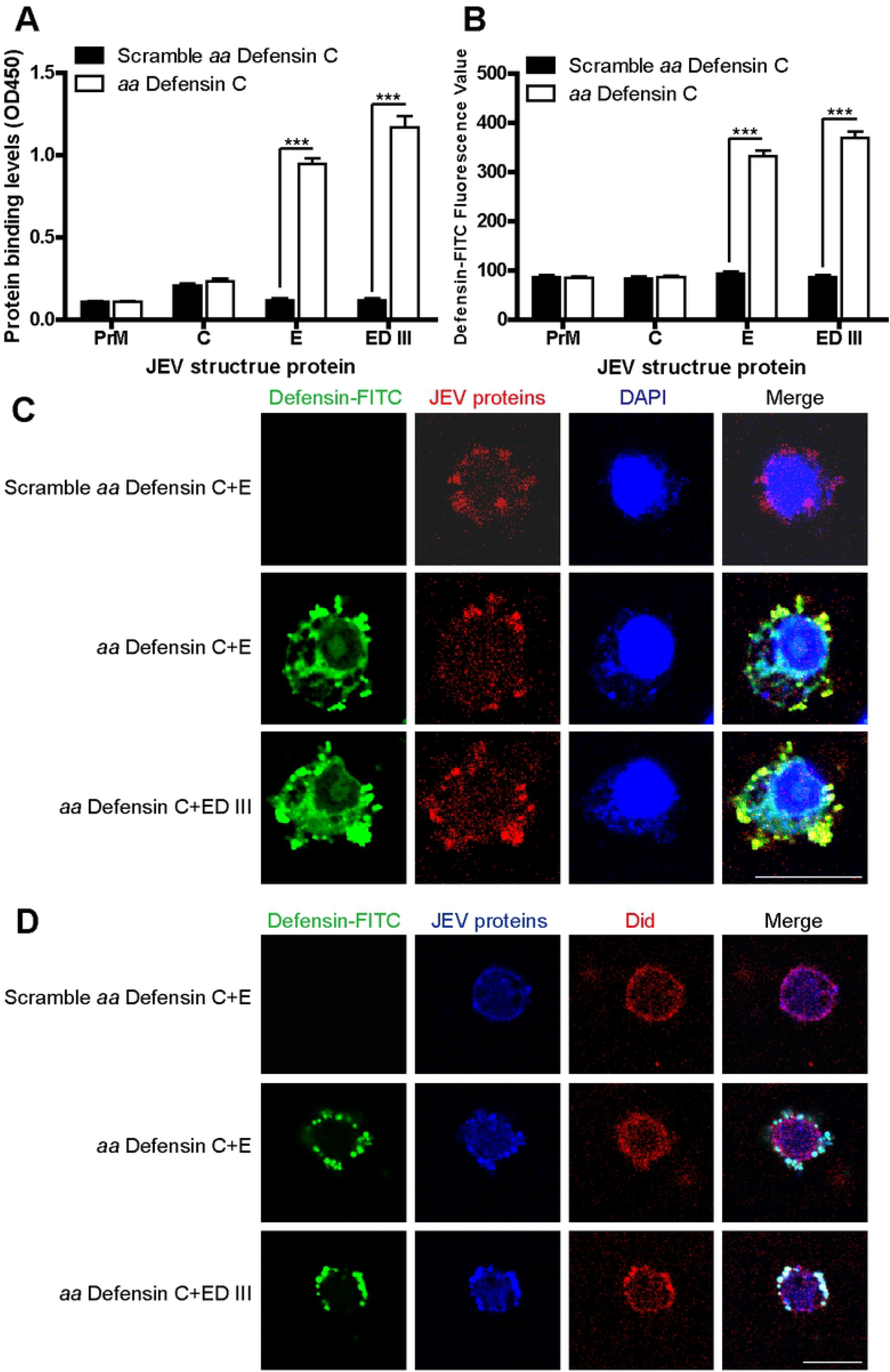
Mosquito defensin bound JEV virions. (A) Viral proteins bind to mosquito defensin. The plate was coated with *aa* defensin C, and incubated with JEV structural proteins. Rabbit polyclonal antibodies to C protein and mouse monoclonal antibody to prM, E and ED III protein were utilized for viral protein binding detection. (B) Mosquito defensin-FITC bind to viral proteins. The plate was coated with purified JEV structural proteins, and incubated with *aa* defensin C-FITC. The fluorescence value of each well was measured. (C and D) Colocalization between defensin and E or ED III proteins. Defensin-FITC and E or ED III were pre-mixed at 4°C and inoculation into C6/36 cells on ice for 4 h. Unabsorbed defensin and proteins were removed by washing with PBS three times. (C) The cells were stained with monoclonal antibody and DAPI to observe JEV E (red fluorescence), defensin-FITC (green fluorescence) and nuclei (blue fluorescence). (D) The cells were stained with monoclonal antibody and Did to observe JEV E (cyan fluorescence), defensin-FITC (green fluorescence) and membrane (red fluorescence). Bar, 10 μm. All experiments were done in triplicate and were performed at least three times. Data are shown as Mean values ± standard deviations.

### LRP2 is responsible for mosquito defensin mediated JEV adsorption

As an extracellular protein, defensin has been reported to interact with receptors on the cell surface and consequently affect intracellular signaling networks. To define the relationship of defensin/cell surface receptor and adsorption enhancement, we analyzed cell-surface receptors that interact with defensin. We knockdowned the expression of a series of potential receptors on the cell surface through RNA interference (RNAi) and found that lipoprotein receptor-related protein 2 (LRP2) responsible for defensin binding [30, 31]. The results indicated that LRP2 interfered with the interaction between defensin-FITC and C6/36 cells (Fig. 5A and Fig. 5B), thus indicating that LRP2 related to the adsorption of extracellular defensin. We further studied the role of LRP2 on JEV adsorption mediated by defensin. Based on significantly RNA knock down (Fig. S3C), No differences were observed between cells with or without LRP2 interference when infected with JEV alone (Fig. 5C). However, when C6/36 cells were incubated with mixed defensin and JEV, a lower JEV level was observed in LRP2 interfered cells. The JEV mRNA level on LRP2 interference cells was 2.8 fold lower than that of the scramble (Negative control, NC) interference group (Fig. 5C), and both the TCID_50_ level and fluorescence value decreased significantly (Fig. 5D and Fig. 5E). In *in vivo* mosquito experiments, the mosquitoes were inoculated with LRP2 or NC siRNA for 3 days (Fig. S3D), and inoculated with defensin and JEV mixture. Whole body samples were collected at 3 days after infection. The JEV mRNA level was significantly lower in LRP2-interference group than in the NC group (Fig. 5F), thus indicating that LRP2 participated in defensin mediated viral adsorption. Additionally, the results of indirect immunofluorescence were in accordance with the above-mentioned results.

**Fig. 5.**
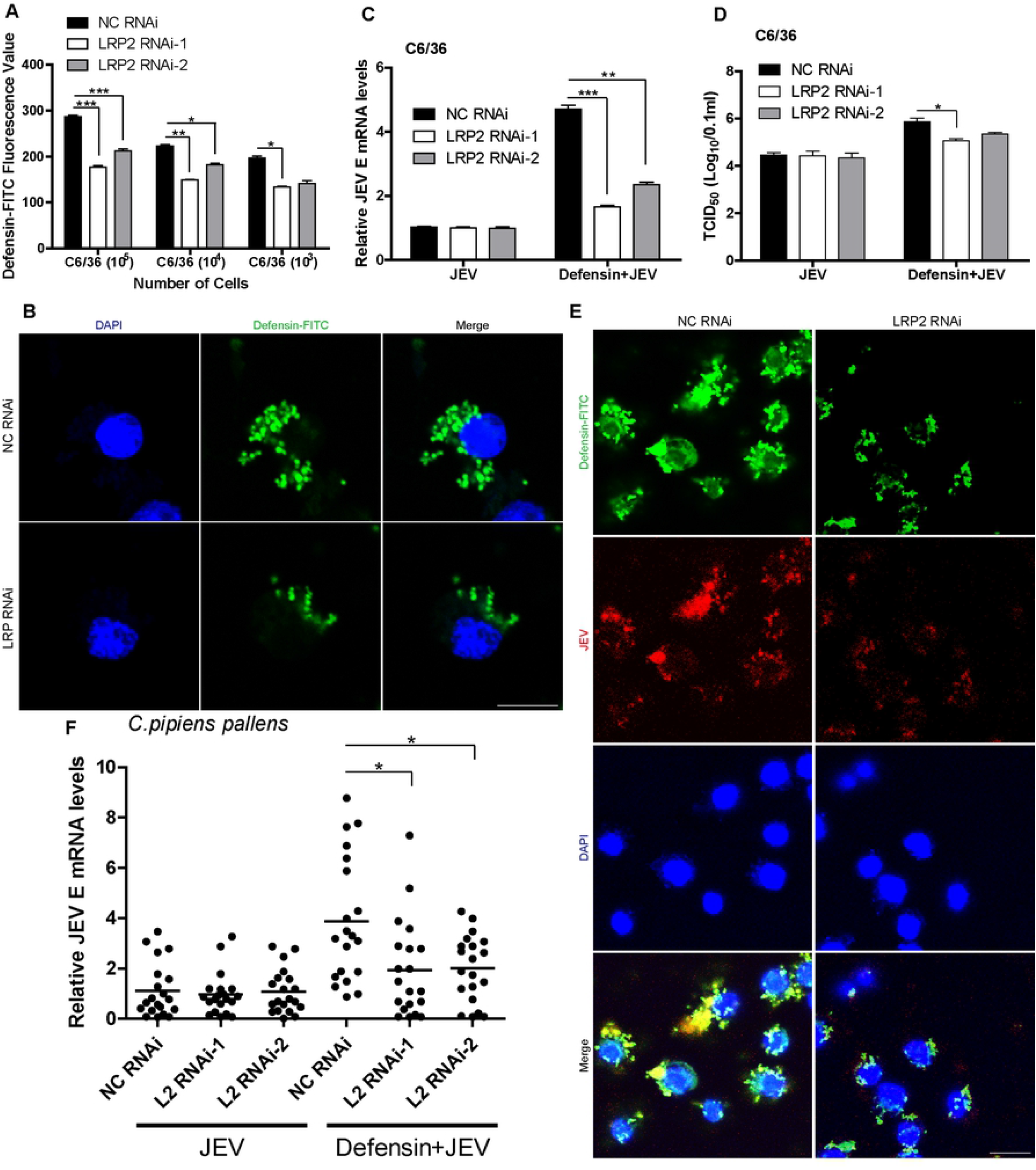
LRP2/defensin pathway mediates JEV adsorption. (A and B) Defensin adsorption was influenced by LRP2. The polylysine treated plate was coated with C6/36, LRP2 siRNAs were transfected into cell. Defensin-FITC was inoculated into cells at 24 h post transfection. After incubation on ice for 2 h, unabsorbed defensin was removed by washing with PBS three times. Fluorescence value was detected by fluorescence analyzer (A) or fluorescence microscope (B). (C, D and E) JEV adsorption on C6/36 cell was influenced by LRP2/defensin pathway. The polylysine treated plate was coated with C6/36, LRP2 siRNAs were transfected into cell. Pre-mixed JEV and *aa* Defensin C was inoculated into cells at 24 h post siRNA transfection. After incubation at room temperature or on ice for 2 h, unabsorbed defensin and virus were removed by washing with PBS three times. For real-time PCR (C) and TCID_50_ (D) measurement, C6/36 cell and supernatant were collected at 2 days post infection. For IFA assay, C6/36 cell was treated immediately after inoculation on ice (E). JEV E was labeled with monoclonal antibody (red fluorescence). Defensin-FITC was detected by green fluorescence, and nuclei were stained with DAPI (blue fluorescence). Bar, 10 μm. (F) *In vivo* JEV adsorption was influenced by LRP2/defensin pathway. Three days after mosquitoes were injected with LRP2 siRNA, the mosquitoes were injected with pre-mixed JEV and defensin. 6 days post infection, mosquitoes were collected to detect JEV E mRNA levels in whole body. All experiments were done in triplicate and were performed at least three times. Data are shown as Mean values ± standard deviations.

Take together, our findings indicated that LRP2 is the cell surface factor responsible for defensin mediated JEV adsorption. LRP2/defensin is a pathway mediates JEV adsorption in mosquito. Lipoprotein receptor-related protein 4 (LRP4) and CXCR4 also showed binding activity with defensin, but this activity did not influence JEV adsorption (Data not shown).

### Mosquito defensin down-regulates the expression of HSC70B on the C6/36 surface and reduces antiviral activity of cell

Defensin can interact with cell-surface receptors and consequently affect signal transduction in cells. Therefore, we employed stable isotope labeling with amino acids in cell culture (SILAC) labeling and mass spectroscopy (MS) methods (Fig. 6A) to determine whether mosquito defensin influences the expression of proteins on cell surface and consequently affects JEV adsorption [32]. Briefly, C6/36 cells were continuously passaged on media with light, medium and heavy stable isotopes. After more than 99.0% cells were labeled with stable isotopes, the cells were then grouped, inoculated with JEV or defensin, and collected according to the plan (Fig. 6A). Cell membrane proteins were extracted for MS analysis. The results showed that HSC70B, a potential mosquito antiviral protein [33], was significantly down-regulated in all defensin, JEV, and defensin + JEV treatments (Fig. 6B). We prepared a rabbit polyclonal antibody against C6/36 HSC70B (UniProt accession number A0A0E3J979) according to the MS results (Fig. S2). Western-blot analysis validated the down-regulation of HSC70B on the C6/36 cell surface in all three treatments (Fig. 6C).

**Fig. 6.**
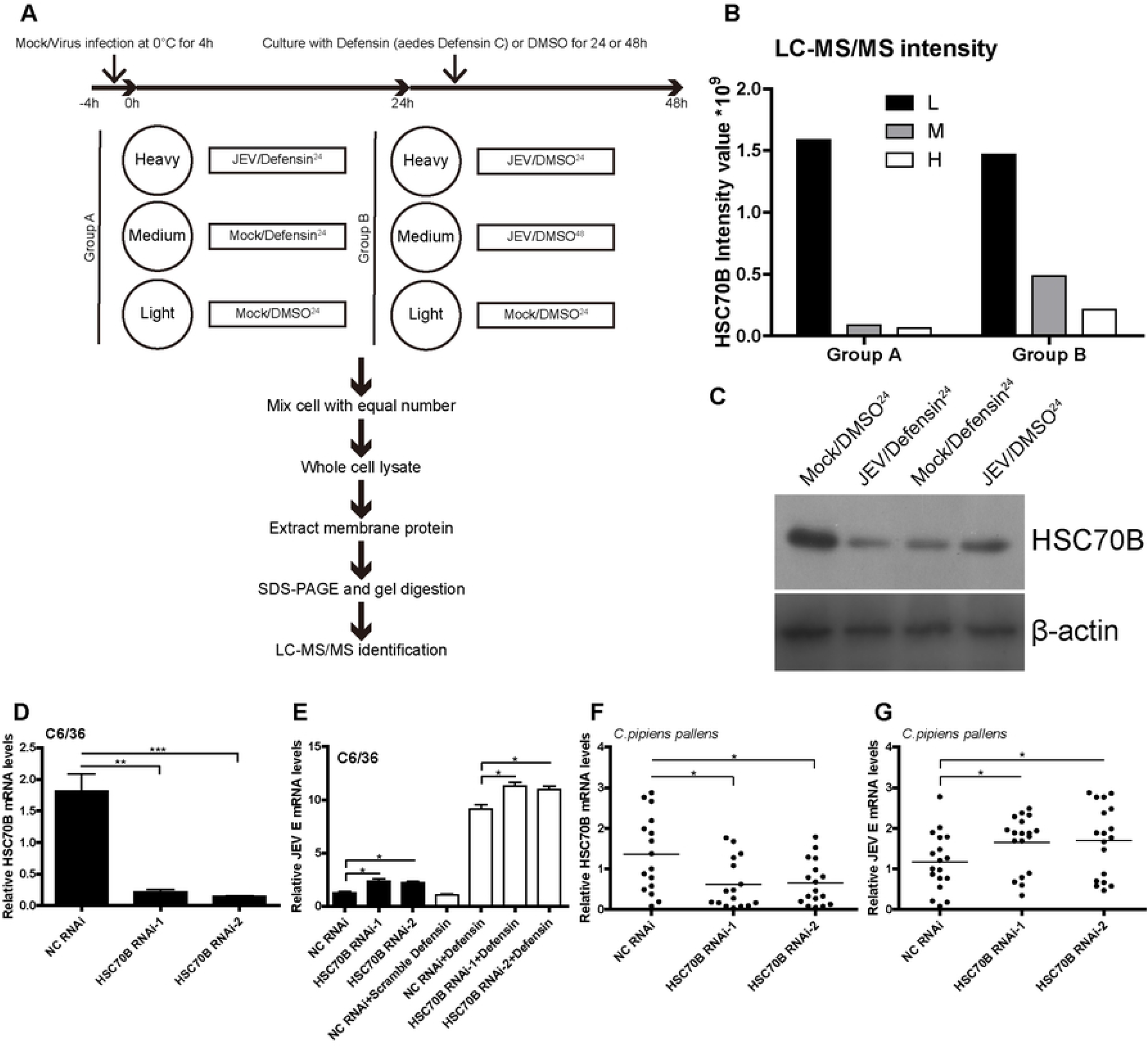
Defensin down-regulated HSC70B on the C6/36 cell surface to enhance JEV adsorption. (A) SILAC/MS workflow. (B) LC-MS/MS intensity of HSC70B on the C6/36 cell surface. Intensity of HSC70B on cell surface was calculate. Protein levels were normalized in a mass spectrometry computing program. (C) Validation of HSC70B expression on C6/36 cell surface according to SILAC/MS. Mosquito HSC70B was probed by rabbit polyclonal anti-HSC70B antibody. (D) The efficiency of HSC70B RNAi *in vitro*. HSC70B siRNA target *aa* HSC70B was transfected into C6/36 cells for 24 h. Cell was collected and HSC70B mRNA was measured by real-time PCR. (E) HSC70B interference facilitated JEV adsorption to cells. C6/36 cells were inoculated with HSC70B siRNA for 24 h and then inoculated with JEV or JEV and defensin on ice for 4 h. Unabsorbed JEV or defensin was removed by washing with PBS three times. The cells were collected to quantify JEV E mRNA levels by real-time PCR. (F) The efficiencies of RNAi HSC70B *in vivo*. siRNA target *cpp* HSC70B was injected. The mosquitoes were collected at 3 days post injection to detect HSC70B mRNA levels *in vivo*. (G) HSC70B interference facilitated JEV infection *in vivo*. Three days after mosquitoes *in vivo* HSC70B RNAi, the mosquitoes were injected with 10 MID_50_ JEV. Mosquito samples were collected at 6 days post infection, and JEV E mRNA levels were detected by real-time PCR. All experiments were done in triplicate and were performed at least three times. Data are shown as Mean values ± standard deviations.

Because HSC70B is a potential mosquito antiviral protein, we tested the function of mosquito HSC70B on JEV adsorption and infection. siRNA targeting C6/36 HSC70B was transfected, and the interference efficiency of HSC70B on C6/36 was detected through real-time PCR analysis (Fig. 6D). Afterward, JEV was inoculated to HSC70B interfered C6/36 cells. The JEV adsorption capacity in HSC70B-interference cells was significantly higher than that in NC-interference cells (*p*<0.05) (Fig. 6E). Additionally, JEV replication level was detected in C6/36 cells treated with HSC70B siRNAs and defensin peptides. Compared to NC group, HSC70B interference significantly heightened the enhancement function of defensin (Fig. 6E). Similarly, we designed siRNA targeting the homologous gene of *cpp* HSC70B. siRNA was injected into *cpp*, and the interference efficiency of HSC70B was detected through real-time PCR (Fig. 6F). JEV mRNA levels were detected at 6 days post infection. Likewise, HSC70B-interfered mosquitoes produced more JEV copies than NC group (Fig. 6G).

Two mechanisms underlying the facilitation effect were identified. One was a direct binding effect, enhancing JEV affinity to the cell surface. The other was an indirect effect, weakening the host defense by down-regulating antiviral HSC70B expression.

### Mosquito defensins facilitate JEV dissemination in salivary gland

To assess the transmission potential of JEV enhanced by mosquito defensins, we detected the virus levels within salivary gland of defensin-treated mosquitoes [34, 35]. Both microinjection and blood meal methods were used in this experiment. JEV and mosquito defensin were mixed before inoculation. Mosquitoes injected with JEV and defensin peptide were collected at 7 or 10 days post infection. Fresh salivary glands were isolated and detected by using real time PCR. JEV level were significantly increased in *Cpp* and *aa* defensin groups (Fig. 7A). JEV level in *Cpp* defensin group indicated 3.5 fold higher at day 7 post infection and 3.1 fold higher at day 10 post infection than that of scramble defensin group in salivary gland. *aa* defensin showed the same role as Cpp defensin did. We further employed blood meal to measure the effect of defensin in JEV dissemination in mosquito salivary gland. Five day-old female Mosquitoes were deprived of sucrose and water 24 h prior to blood meal. Mosquitoes were then feed with infectious blood with JEV and defensin peptides for 2 h. Fresh blood was collected from health mice and delivered through Hemotek membrane feeding apparatus. 2ml blood with JEV (5×10^6^ TCID_50_) and peptide (200 μM) were used for each groups. JEV level in *Cpp* defensin group showed 4.2 fold higher at day 7 and 10 post infection than that of scramble defensin group in salivary gland (Fig. 7B). JEV level in *aa* defensin group also showed higher results than scramble group. These results implied that mosquito defensins facilitate JEV dissemination in mosquitoes and increase transmission potential after infection.

**Fig. 7.**
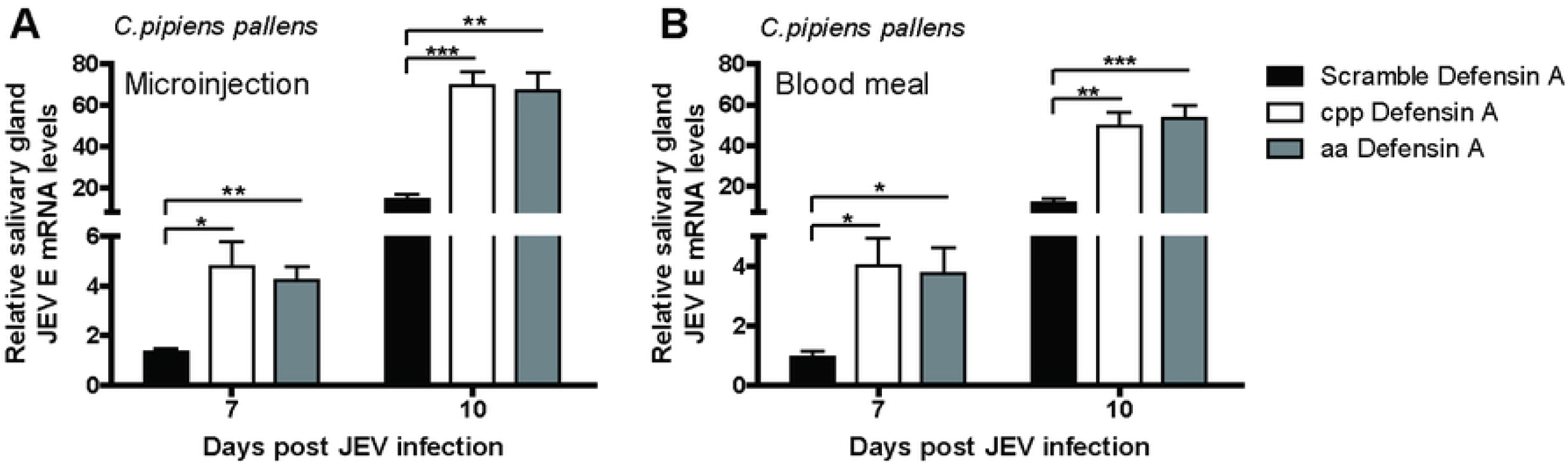
Mosquito defensin enhanced JEV replication in salivary gland. (A) JEV E mRNA levels within salivary gland based on microinjection. *Cpp* defensin A (100 μM) and JEV (10 MID_50_) were pre-mixed at 4°C for 2 h and injected into female mosquitoes. Salivary glands were isolated at 7 and 10 days post injection and detected by real-time PCR. (B) JEV E mRNA levels within salivary gland based on blood meal. *Cpp* defensin A (100 μM) and JEV (10^3^ MID_50_) were pre-mixed at 4°C. Mixture was added into fresh blood with anticoagulant. Blood meal was performed for 2 h. Salivary glands were isolated at 7 and 10 days post infection and detected by real-time PCR. All experiments were done in triplicate and were performed at least three times. Data are shown as Mean values ± standard deviations.

## Discussion

JEV is a serious mosquito borne disease common in Asia-Pacific tropical and subtropical regions [2, 9, 10]. More than 100,000 people are at risk. Moreover, JEV can cause death or permanent sequelae. Pigs are the reservoir host of the virus. Humans, horses and other animals are dead-end hosts. Mosquitoes, especially *culex*, are the most important vector [4]. At present, the prevention and control of JEV mainly relies on vaccine immunization, whose protection time is limited. JEV remains a threat to health and even life for immunocompromised children or older people [6, 7]. With the increasing problem of global warming, the clinical incidence of JEV is increasing [36]. Few mechanistic studies have focused on the JEV transmission by mosquito vectors. It is of practical significance to understand the interaction between JEV and mosquito vectors and the immune escape mode of JEV in controlling this mosquito borne disease.

In this study, we analyzed the gene expression of defensin from *Cpp* and *aa*. Defensin A and an unnamed defensin from *Cpp*, defensin A, B and C from *aa* shared high sequence similarity, thus indicating similar functions of these defensins. Subsequently, we confirmed that defensin A and defensin C are the main defensin types of *cpp* and *aa*, respectively. Given the high similarity of the amino acid sequences, we synthesized only *cpp* defensin A and *aa* defensin C in further studies. The nucleotide sequence of *cpp* defensin A (number MH756645) has been submitted to the NCBI database.

The up-regulation of defensin after JEV infection was consistent with reports on other flavivirus viruses [37]. The highest up-regulation was observed at 7 days post infection. From the organism perspective, the defensin in the salivary gland and whole body was up-regulated more than that in the midgut. JEV replication in salivary gland, the most sensitive tissue to JEV [18], was positively correlated with defensin level.

The mature peptide of defensin was utilized to study the role of mosquito defensin in JEV infection [25, 38]. In general, defensin is fewer than 60 amino acids and is processed from a precursor protein. In this study, mosquito defensin and human defensin β2 were composed of 40 amino acids and 34 amino acids [39], respectively. Only 11% sequence similarity was identified between mosquito and human defensin. Unexpectedly, mosquito defensin facilitated JEV infection, in contrast to human defensin, but the facilitation effect was exerted only on mosquito cells or mosquitoes. Thus, JEV utilized the host defense system, reflecting its “intelligence” in infection [40]. However, mosquito defensin inhibited JEV infection in mammalian cells, thus indicating its varied mechanisms of action and the complicated interaction between virus and host [19].

Further analysis demonstrated that mosquito defensin facilitated JEV adsorption to target cells by directly binding JEV virions [27, 41]. By screening JEV structural proteins, we found that mosquito defensin bound the ED III domain of JEV E. ED III is a crucial domain of JEV that is responsible for the production of neutralizing antibody [42]. The antiviral effect of mosquito defensin on JEV is likely to be due to its binding the ED III domain and subsequent virion destruction [19, 25]. Because mosquito defensin facilitated JEV infection, the binding of defensin and ED III can be inferred to have only weak antiviral effects. Nevertheless, this binding enhanced virion adsorption ability to a large extent. The broad transmission of JEV by mosquitoes is ascribed to both the crude immune system of mosquitoes and the infection strategy of the virus. Mosquito defensin could improve the adsorption ability of JEV on target cells. Additionally, ELISA results showed that high concentration of mosquito defensin interacted with the target cells without the assistance of viruses.

Defensin receptors expressed on the cell surface may lead to enhanced adsorption. We scanned the potential cell-surface receptor proteins of defensin through RNAi and found that the LRP2-defensin pathway was responsible for JEV adsorption. In mammalian animals, LRP2 is the receptor for defensin, regulating the contraction of smooth muscle cells by combining with human alpha defensin [30, 31]. However, the roles of LRP2 in mosquitoes have not been reported. In the present study, we demonstrated that LRP2 participates in JEV adsorption mediated by defensin. JEV first binds defensin, and then, owing to the affinity of defensin for LRP2, the defensin/JEV complex adsorbs to the cell surface more readily, thereby increasing the chance of infection. This proposed mechanism of promotion of JEV infection by defensin/LRP2 is similar to that for JEV and WNV mediated by C type lectin/PTP-1. That is, virus first combines with extracellular secrete proteins with high affinity to cells, and this is followed by binding to cell surface receptor to infect target cells.

Given that mosquito defensin directly interacts with mosquito cell surface receptors, we analyzed how it regulates cell surface proteins. The changes in cell surface proteins were determined through SILAC and MS analysis [32]. We identified a potential antiviral protein, HSC70B, that is significantly down-regulated by defensin or JEV treatment [33]. HSC70B inhibited JEV adsorption, as demonstrated through an RNAi approach, thus indicating that mosquito defensin indirectly affects JEV adsorption by regulating cell surface antiviral protein expression. However, this indirect effect was found to be lower than the direct defensin binding effect. Together, our findings indicated that the effect of mosquito defensin on JEV is composed of weak antiviral effect, direct binding enhancement and indirect immune regulation. Curiously, both defensin and HSC70B are antiviral proteins in mosquito, but it looks like they could’t work together on JEV infection. We did not identify the mechanisms through which defensin down-regulates HSC70B, because of the limited information available on the relevant signal pathways. We deduced that there is a negative feedback mechanism between HSC70B and defensin [43], thus implying that an increase in defensin would decrease HSC70B level. Another possibility may be that HSC70B has varying functions in different conditions, except for the antiviral effect.

JEV infection up-regulated mosquito defensin expression in the salivary gland and defensin also heightened the JEV dissemination in salivary gland, thus suggesting that the defensin may be influence the transmission of JEV by mosquito. Further research on mosquito defensin in JEV cross-species transmission is needed.

To our knowledge, this is the first report on the effects of mosquito defensin on JEV infection in mosquito vectors, revealing a new immune escape mechanism of JEV infection and transmission. This study broadens our knowledge of transmission of JEV as well as other mosquito borne viruses, providing novel insights into viral transmission mechanisms.

## Materials and methods

### Ethics statement

All animal experiments were performed in compliance with the Guidelines on the Humane Treatment of Laboratory Animals (Ministry of Science and Technology of the People’s Republic of China, Policy No. 2006 398) and were approved by the Institutional Animal Care and Use Committee at the Shanghai Veterinary Research Institute (IACUC No: Shvri-Pi-0124).

### Cells, defensin and viruses

Baby hamster kidney (BHK-21) and African green monkey kidney (Vero) cells were purchased from the ATCC (Rockville, Maryland) and maintained in Dulbecco’s modified Eagle’s medium (DMEM) supplemented with 10% FBS at 37°C in a 5% CO_2_ incubator. C6/36 cells (ATCC) were cultured in RPMI-1640 medium supplemented with 10% FBS at 28°C.

Mature *Cpp* defensin A (NCBI accession number: MH756645), *aa* defensin C (NCBI accession number: XP_019527114.1), human defensin β2 (NCBI accession number: NP_004933.1) and scrambled defensin peptides (purity ⩾ 99%) were synthesized by WC-Gene Biotech Ltd. (Shanghai, China). The amino acid sequences are shown in Table S3. The defensins were dissolved in DMSO (for cell, *ex vivo* or *in vivo* experiments) or PBS (for ELISA detection) and stored at room temperature. Defensins labeled with FITC were kept in the dark at room temperature.

JEV strain N28 (NCBI accession number: GU253951.1) was stored in our laboratory and propagated in C6/36 cells. The TCID_50_ and MID_50_ of the virus were measured in BHK-21 cells or female mosquitoes and calculated by the Reed-Muench method [17, 44].

### Infection and RNA interference *in vitro*

Defensins or scrambled defensin peptides were pre-mixed with JEV (MOI=0.1) at 4°C, then inoculated into cells. C6/36 cells were incubated at 28°C for 2 h. Vero and BHK-21 cells were incubated at 37°C for 2 h. At 24–120 h post infection, the supernatant or cells were collected. Viral titer was determined by TCID_50_ method and mRNA expression levels were measured by real-time PCR. To determine JEV adsorption, defensins were pre-mixed with JEV at 4°C for 2 h. C6/36 cells were incubated with the mixture on ice for different times. Unabsorbed JEV was removed by washing with PBS for three times. The cells were collected for JEV E mRNA quantification or other measurements.

For the *in vitro* RNA interference, siRNA (Table S1) was transfected into C6/36 cells with Cellfectin II reagent (Invitrogen). JEV was inoculated at 24 h post transfection. At 72 h post infection, the cells were collected. Total RNA was isolated, and the viral or gene load was determined by real-time PCR.

### Infection and RNA interference *in vivo*

For *in vivo* experiments, 10-fold serial dilutions were made from a 10^9.3^ TCID_50_ JEV stock. Cold-anesthetized 5 day old female mosquitoes were randomly divided into various groups (n≥13). Both microinjection and blood meal methods were carried out in infection experiment. For microinjection, the mosquitoes were infected by microinjection (250 nL) into the thorax. An Eppendorf CellTram oil microinjector and 15 μm needles were used for injecting the mosquitoes. Control mosquitoes were injected with an equivalent volume of PBS [17, 18, 45]. The mosquitoes were harvested, and the viral loading was quantified. For blood meal, fresh blood of specific pathogen free mouse was collected in tubes with anticoagulant. Virus or defensin peptides were mixed and added into fresh blood before feeding. 2 ml blood was used in blood meal by Hemotek FU1 Feeder for each group [34, 46].

*In vivo* RNAi was performed as described previously [18]. The siRNA targeting the *Cpp* genes was synthesized by Genepharma (Shanghai, China). The sequences are shown in Table S1. For RNAi and virus challenge, female mosquitoes at 5 days after eclosion were injected into the thorax with 2 μg dsRNA in 250 nL PBS. After a 3 day recovery period, the mosquitoes were microinjected with JEV at different MID_50_ in 250 nL PBS for functional studies.

### RNA isolation and real-time PCR

For real-time PCR, RNA was extracted from cell suspensions or mosquito samples by Qiagen total RNA isolation kit according to the manufacturer’s instructions. The RNA concentration was measured by NanoDrop spectrophotometer. cDNA was generated by RT Master reverse transcription kit (Takara) according to the manufacturer’s instructions. Real-time quantitative PCR experiments were performed in ABI Prism 7500 sequence-detection system (Applied Biosystems, Foster City, CA) with SYBR Green PCR Master Mix (Takara) according to the manufacturer’s instructions. The primer sequences are listed in Table 1. The thermal cycling conditions were as follows: 10 min at 95°C, followed by 40 cycles of 95°C for 5 s and 60°C for 1 min. All experiments were performed in triplicate, and gene expression levels are presented relative to those of β-actin. The fold change in relative gene expression compared with the control was determined with the standard 2^-ΔΔCt^ method.

### Virus titer

Supernatants were harvested from cell cultures for TCID_50_ assays as described previously [18]. Briefly, BHK-21 cells were seeded on a 96-well plate and grown to 60% confluence. The supernatants were diluted in a 10-fold dilution series and added to each well of the 96-well plate. One hundred microliters of each dilution was added in eight replicate wells, and eight replicate mock controls were set. The plates were incubated at 37°C for 1.5 h. Then the supernatants were discarded and replaced with 100 μL of DMEM supplemented with 1% FBS. After 5 days in culture, the cytopathic effect was recorded. The TCID_50_ of the virus was calculated by Reed-Muench method [44].

### Indirect immunofluorescence and western blotting

Indirect immunofluorescence and western blotting were performed as described previously [18]. The antibodies used were mouse anti JEV E monoclonal antibody, rabbit anti mosquito β actin polyclonal antibody, rabbit anti mosquito HSC70B polyclonal antibody, goat anti-mouse IgG-HRP antibody (1:10000; Santa Cruz), Alexa Fluor 405-conjugated anti-mouse IgG antibody (1:500; Abcam), Alexa Fluor 488-conjugated anti-rabbit IgG antibody (1:500; Thermo Fisher Scientific) and Alexa Fluor 594-conjugated anti-rabbit IgG antibody (1:500; Thermo Fisher Scientific). DAPI and Did were used for nucleus and membrane staining. Immunofluorescence was imaged with a Nikon C1Si confocal laser scanning microscope.

For tissue immunofluorescence assays, salivary glands were isolated on sialylated slides, washed with PBS, fixed with 4% paraformaldehyde for 1 h, and blocked in PBS with 2% bovine serum albumin (BSA) at room temperature for 2 h. The samples were incubated with mixture of JEV and *cpp* defensin A-FITC, detected with mouse anti JEV E monoclonal antibody and imaged with a Nikon C1Si confocal laser scanning microscope.

### Protein expression and ELISA

The purified JEV structural proteins (C, M, E, ED III) from the S2 insect expression system (Invitrogen) were quantified by using the bicinchoninic acid (BCA) assay. Expressed proteins were used for ELISA or IFA analysis [47].

For defensin ELISA, defensin peptide was dissolved in PBS, then was diluted with 0.1 M dicarbonate (pH 9.6) to a final concentration of 250–750 ng. The plate was coated overnight and incubated with 2% BSA for 2 h. Afterward, 100 μL JEV virus (1×10^5^ TCID_50_) was added and incubated for 30 min at room temperature. The wells were washed with PBST five times, mouse polyclonal antibody to JEV was added to the wells and incubated for 30 min. The wells were washed with PBST five times, and goat anti-mouse antibody labeled with HRP was added. After incubation at room temperature 30 min and washing with PBST five times, TMB was added to the wells as a chromogenic substrate. The plate was developed in the dark for 10 min, and H2SO4 was added to stop the reaction. The absorbance of each well was read at 450 nm.

For viral protein ELISA, purified JEV structural proteins diluted in 0.1 M dicarbonate (pH 9.6) were added to the plate wells. The plate was coated overnight and incubated with 2% BSA for 2 h. Then 100 μL defensin (50 μM) labeled with FITC was added. The plate was incubated for 30 min at room temperature and washed with PBST five times before fluorescence measurement.

For C6/36 cell ELISA, the plates were pre-treated with polylysine. Healthy and fresh C6/36 cells were counted and diluted with 0.1 M dicarbonate (pH 9.6) to a final concentration of 1×10^5^ cells per well. The plate was processed as described above for JEV structural proteins or defensin coated ELISA.

### SILAC/MS analysis

C6/36 cells were continuously passaged for eight generations on media with light, medium and heavy isotopes. All three labeling efficiencies reached 99%. The cells were grouped, inoculated with JEV or defensin, and collected at 24 h or 48 h according to the procedure. Equal amounts of cells from light, medium and heavy media in the same group were mixed to extract cell membrane proteins according to the manufacturer’s instructions (Pierce). Extracted membrane proteins were quantified by BCA, identified by MS and normalized for further analysis.

### Statistical analysis

All experiments were carried out in at least triplicate. Mean values ± standard deviation (SD) were calculated in Microsoft Excel. Statistical analysis was done with Student’s t tests, and values were considered significant when *p*<0.05. Figures were created in GraphPad™ Prism 5.0 software.

## Acknowledgments

This work was supported by the the National Key Research and Development Program of China (No. 2017YFD0501805, 2018YFD0500101), the Shanghai Natural Science foundation (no. 18ZR1448900), applied research on disease prevention and controls in military (No. 13BJYZ27), the National S & T Major Program (No. 2012ZX10004-220).

## Supplement Figures

**Fig. S1. Sequence and abundance of mosquito defensins.** (A) Amplification of *cpp* defensin A by PCR. (B) Defensin sequence alignment. Alignment of *cpp* defensin sequences (*Cpp* defensin A and unnamed defensin). (C) Abundance of defensins in C6/36 and *cpp*. Defensin genes were amplified and cloned into pMD18 plasmids, and positive plasmids were used to construct standard curves. Defensin abundance in cells or mosquitoes was quantified with a standard curve through real-time PCR. Defensin abundance is shown as a proportion. Target defensins are shown in gray in columns; the total column represents total defensins. (D, E) Defensin sequence alignment. Alignment of defensins in different mosquito Species (D). Alignment of mosquito defensins and human defensin (E). Alignment was performed by DNAMAN software. Data are shown as Mean values ± standard deviations.

**Fig. S2. Major sequences used in this study.** (A) *Culex pipiens pallens* defensin A protein sequence (NCBI number MH756645); the mature defensin sequence is in red. (B) *Culex pipiens pallens* HSC70B partial sequence. (C) *Culex pipiens pallens* unnamed defensin partial sequence. (D) C6/36 HSC70B protein sequence (immunogenic peptide for antibody preparation is in red).

**Fig. S3. RNA interference efficiency in *in vitro* and *in vivo***. (A) The efficiency of defensins RNAi *in vivo*. siRNAs target *cpp* total defensins or defensin A were injected. Mosquitoes were collected at 3 days post injection to detect defensins mRNA levels by real-time PCR. (B) The efficiency of defensins RNAi *in vitro*. siRNAs target *aa* defensins was transfected into C6/36 cells for 24 h. Cell was collected and defensins mRNA were measured by real-time PCR. (C) The efficiency of LRP2 RNAi *in vitro*. LRP2 siRNA target *aa* LRP2 was transfected into C6/36 cells for 24 h. Cell was collected and LRP2 mRNA was measured by real-time PCR. (D) The efficiency of LRP2 RNAi *in vivo*. siRNA target *cpp* LRP2 was injected. Mosquitoes were collected at 3 days post injection to detect LRP2 mRNA levels by real-time PCR. All experiments were done in triplicate and were performed for three times. Data are shown as Mean values ± standard deviations.

